# Primary tumor transcriptomic signature is associated with locoregional dissemination in canine mammary carcinoma

**DOI:** 10.64898/2026.09.10.750581

**Authors:** Jin-Hyuk Park, Jun-Yeol Choi, Se-Hoon Kim, Ki-Hoon Song, Seon-Kyu Kim, Kyoung-Won Seo

## Abstract

Canine mammary carcinoma (CMC) exhibits heterogeneous metastatic behavior, while accurate assessment of locoregional dissemination may be limited by incomplete sampling, occult metastasis, and variable sentinel lymph node anatomy. We investigated whether primary-tumor transcriptomic programs are associated with locoregional dissemination. The training cohort comprised 113 CMC RNA sequencing samples classified by peritumoral lymphatic vessel invasion, and the independent evaluation cohort comprised 27 microarray samples classified by histopathologically confirmed lymph node (LN) metastasis. Expression profiles were mapped to human ortholog gene symbols, and single-sample gene set enrichment analysis (ssGSEA) was performed using Molecular Signatures Database (MSigDB) gene sets. Seven candidate gene sets selected through univariable screening and literature-guided curation were entered into least absolute shrinkage and selection operator (LASSO) logistic regression, yielding a four-gene-set transcriptomic risk score (RS). The RS showed an apparent receiver operating characteristic area under the curve (ROC-AUC) of 0.949 (95% confidence interval, 0.906–0.993) and precision-recall area under the curve (PR-AUC) of 0.725 in the training cohort; bootstrap internal validation yielded an optimism-corrected ROC-AUC of 0.920. In the evaluation cohort, the ROC-AUC was 0.890 (95% confidence interval, 0.766– 1.000) and PR-AUC was 0.869. Discrimination was attenuated but retained in histologically restricted sensitivity analyses, and higher RS remained associated with locoregional dissemination after clinicopathologic adjustment. These findings support gene-set-level transcriptomic scoring as a potential complementary approach for characterizing dissemination-associated primary-tumor states in CMC.

## Introduction

Canine mammary carcinoma (CMC) is a common malignant tumor in female dogs [1,2] and exhibits marked heterogeneity in histologic subtype and clinical behavior [3–6]. Regional lymph nodes and the lungs have been reported as common sites of metastasis in CMC [4]. Lymphatic vessel invasion [7,8] and regional lymph node metastasis [8–11] are important prognostic factors associated with adverse outcomes. Therefore, the assessment of locoregional dissemination is important for staging [12] and may inform subsequent treatment planning [6].

Histopathologic evaluation of excised regional lymph nodes remains the definitive method for confirming nodal metastasis [13]. However, nodal staging in CMC has several limitations. Although regional lymph node evaluation or resection has traditionally been guided by tumor location [13], anatomically regional lymph nodes may not always correspond to the true sentinel lymph nodes because of the variable lymphatic drainage patterns [14]. Sentinel lymph node mapping may improve the identification of the true draining lymph nodes [15,16], but this approach is not yet standardized or routinely implemented in veterinary oncology practice [17]. Even after lymph node excision, small metastatic foci such as micrometastases or isolated tumor cells may be missed by routine hematoxylin and eosin (H&E)-based histopathologic evaluation [18,19].

Gene-expression studies in human cancer have suggested that metastatic behavior may be partly encoded in the transcriptional state of the primary tumor. In breast cancer, primary-tumor expression profiles have been used to identify signatures associated with subsequent distant metastasis, including the 70-gene signature reported by van’t Veer et al. [20] and the 76-gene signature reported by Wang et al. [21]. More broadly, Ramaswamy et al. identified a metastasis-associated transcriptional signature across solid tumors that was also detectable in subsets of primary tumors, supporting the concept that metastatic propensity may be reflected in primary-tumor gene-expression programs [22].

In veterinary oncology, molecular features associated with metastatic behavior in CMC have been investigated at multiple levels. Transcriptome-wide profiling has demonstrated distinct gene-expression patterns in primary tumors with lymph-node metastasis [23], while subsequent studies have identified metastasis-associated microRNAs (miRNAs) [24] and individual transcripts related to subsequent metastatic progression and survival [25]. However, most reported markers have remained individual candidates or cohort-specific signatures, and their reproducibility across independent populations has been limited. For example, a three-transcript signature derived from a canine cohort with nodal metastasis did not discriminate lymphatic invasion in an independent RNA sequencing (RNA-seq) cohort, with only one component, *SFRP1*, showing partial replication in subsequent tissue cohorts [26].

Together, these findings suggest that metastatic behavior is reflected in the primary-tumor transcriptome, while also highlighting the difficulty of transferring individual transcript-based markers across cohorts and transcriptomic platforms. Because locoregional dissemination is likely to involve coordinated biological processes rather than isolated gene-level alterations [27,28], evaluating transcriptional activity at the gene-set level may provide a complementary approach for capturing dissemination-associated transcriptional features across cohorts. Single-sample gene set enrichment analysis (ssGSEA), implemented within the gene set variation analysis (GSVA) framework, enables sample-wise quantification of predefined gene-set activity [29,30] and may improve comparability between RNA-seq and microarray datasets [31].

In this study, we aimed to develop a transcriptomic risk score (RS) associated with lymphatic spread by applying the least absolute shrinkage and selection operator (LASSO) logistic regression to ssGSEA-derived gene-set scores from primary CMC transcriptomes. We then evaluated the RS in an independent microarray cohort using histopathologically confirmed lymph node (LN) metastasis as an indicator of locoregional dissemination. The resulting RS was associated with lymphatic spread in the training cohort and retained discriminatory ability for LN metastasis status in the independent evaluation cohort.

## Methods

### Study cohorts and datasets

We retrieved publicly available RNA-seq and microarray datasets from the Gene Expression Omnibus (GEO). The training cohort was derived from the RNA-seq dataset of canine mammary gland tumors reported by Kim et al. (GSE119810). Of the 157 RNA-seq tumor samples available in the dataset, 36 benign mammary tumors and 8 malignant tumors with sarcomatous, carcinosarcomatous, or predominantly spindle-cell histologies (one fibrosarcoma, three osteosarcomas, three carcinosarcomas, and one spindle cell carcinoma) were excluded, leaving 113 mammary carcinomas for the training cohort [32]. The evaluation cohort consisted of microarray data from 27 CMC cases reported by Klopfleisch et al. (GSE20718) [23], including 13 lymph node (LN)-positive and 14 LN-negative tumors. All tumors were histologically classified as simple carcinomas and characterized by invasive and predominantly solid growth patterns. The original study deliberately selected the LN-negative group – using primary-tumor size as one selection criterion – to represent invasive malignant tumors without nodal metastasis, rather than a biologically indolent comparator group. Clinicopathologic variables used in the present analyses, including histologic grade, were obtained from the case-level GEO metadata. Clinical T stage, which was not directly annotated in the GEO metadata, was derived from the recorded primary-tumor diameter according to the adopted tumor-node-metastasis (TNM) criteria (T1, < 3 cm; T2, 3–5 cm; T3, > 5 cm) [33,34].

### Outcome definition

Histologic annotations for the training cohort were obtained from the original study, in which all tumors were evaluated by two pathologists. Cases with tumor cells identified within peritumoral lymphatic vessels were classified as Spread-positive, whereas cases without evidence of lymphatic invasion or nodal metastasis were classified as Spread-negative. For the evaluation cohort, nodal metastasis status was likewise obtained from the original study, in which lymph node histology had been assessed by two board-certified pathologists. This histologically defined nodal status was used to evaluate the association between the training-derived transcriptomic risk score and locoregional dissemination.

### Ethics statement

This study was a secondary analysis of publicly available datasets and involved no new animal enrollment, intervention, sample collection, or experimentation. Therefore, institutional animal ethics approval was not required for the present study.

### Data preprocessing

For the 113 CMC RNA-seq samples, transcript-level abundance estimates were generated using Salmon (version 1.10.2) [35] and summarized to gene-level transcripts per million (TPM) values using tximport (version 1.34.0) [36] with a transcript-to-gene (tx2gene) mapping based on Ensembl release 114 of the ROS_Cfam_1.0 [37]. Gene-level expression values were transformed as *log_2_(TPM+1)* for downstream ssGSEA. We mapped canine Ensembl gene IDs to human ortholog gene symbols using Ensembl orthology information retrieved through the biomaRt R package (version 2.62.1) to enable scoring against the human-derived Molecular Signatures Database (MSigDB) gene sets. When multiple canine genes mapped to the same human ortholog gene symbol, the expression values were collapsed by taking the median.

For the evaluation cohort, gcRMA-preprocessed microarray expression data were downloaded from GEO [23]. Probe IDs were mapped to canine Ensembl gene IDs using the canine2.db annotation package (version 3.13.0), followed by canine-to-human ortholog mapping using the same Ensembl/biomaRt procedure described above [37,38]. Probes without mapped human ortholog gene symbols were excluded. When multiple probes were mapped to the same human ortholog gene symbol, expression values were collapsed by taking the median across probes.

### Feature selection and model development

We obtained gene sets from MSigDB v2025.1.Hs, including the Hallmark (H), C2 curated, and C5 ontology collections [39–41]. To ensure consistent gene-set membership across the RNA-seq training and microarray evaluation cohorts, each gene set was restricted to human ortholog genes represented in both datasets. ssGSEA was then performed separately in each cohort using the GSVA R package [29,30] with normalization disabled (normalize = FALSE). The resulting scores were z-standardized independently within each cohort before downstream analyses.

### Gene-set screening using univariable logistic regression

For each gene set, we performed univariable logistic regression in the training cohort using lymphatic spread status as the binary outcome and the z-standardized ssGSEA score as the predictor. *P* values from all 17,675 univariable tests (50 H, 6,357 C2, and 11,268 C5 gene sets) were adjusted together using a single Benjamini–Hochberg (BH) procedure to control the false discovery rate (FDR) [42]. Gene sets with BH-adjusted *P* < 0.05 were retained for literature-guided curation. Of the 17,675 gene sets screened, 2,391 met this threshold, comprising 11 of 50 H gene sets, 1,410 of 6,357 C2 gene sets, and 970 of 11,268 C5 gene sets (S1 Table). To obtain a biologically interpretable and distinct set of candidate predictors, we manually curated gene sets into biological domains relevant to metastatic dissemination and tumor malignancy, including metastasis-related expression programs, proliferation/cell-cycle activity, telomere/genome stability, dedifferentiation, vascular remodeling/permeability, hypoxia, and immune evasion. When multiple gene sets represented overlapping biological processes, a representative gene set was selected based on relevance to mammary tumor biology or metastatic dissemination, preference for well-curated gene-set definitions, and biological interpretability. This procedure yielded seven candidate gene sets for subsequent LASSO logistic regression (S2 and S3 Tables).

### LASSO logistic regression for candidate gene sets

We performed LASSO logistic regression using the glmnet R package with a binomial family and an alpha value of 1 [43,44]. Given the limited number of Spread-positive cases (n = 14), penalty parameter selection was stabilized by performing 100 repetitions of five-fold cross-validation. For penalty selection in the original training cohort, each fold was constrained to include at least two Spread-positive samples. For each repetition, the one-standard-error penalty parameter (λ1SE), defined as the largest λ value with cross-validated binomial deviance within one standard error of the minimum, was recorded (S5 Table). The final penalty parameter was defined as the median λ1SE across the 100 repetitions, and the final model was refitted in the full training cohort using this penalty. The correlation structure among the candidate gene sets was examined using Spearman’s rank correlation analysis (S1 Fig).

To assess optimism in the apparent training performance and the stability of gene-set selection, bootstrap internal validation was performed using 1,000 ordinary bootstrap resamples of the training cohort. Within each bootstrap resample, the penalty-selection procedure was repeated using 100 repetitions of stratified five-fold cross-validation without imposing a minimum number of Spread-positive samples per fold, because the number of positive cases varied across bootstrap resamples. The median λ1SE was then used to fit a LASSO model. The fitted model was evaluated both in the bootstrap sample and in the original training cohort. Optimism for each resample was calculated as the difference between the area under the receiver operating characteristic curve (ROC-AUC) in the bootstrap sample and that obtained when the same model was applied to the original cohort. The mean optimism across 1,000 resamples was subtracted from the apparent ROC-AUC of the final model to obtain an optimism-corrected ROC-AUC. Gene-set selection stability was summarized as the proportion of bootstrap resamples in which each of the seven candidate gene sets had a non-zero coefficient. The bootstrap validation was conditional on the seven candidate gene sets identified through the preceding screening and literature-guided curation procedure.

### Statistical evaluation

We defined the RS as the linear predictor derived from the final LASSO logistic regression model:

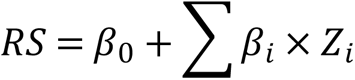

(where β_0_ denotes the intercept, β_i_ denotes the coefficient for each selected gene set, and Z_i_ denotes the z-standardized ssGSEA score for the corresponding gene set.)

Group comparisons were performed using the two-sided Wilcoxon rank-sum test. Sensitivity analyses were additionally performed to assess whether differences in RS according to lymphatic spread status were retained within more histologically comparable subsets. In the training cohort, analyses were repeated after restriction to simple carcinomas, grade 2–3 tumors, and grade 3 tumors; in the evaluation cohort, the comparison was repeated among grade 3 tumors. *P* values for the three subgroup analyses in the training cohort were adjusted using the BH method. Apparent discrimination of the final RS in the full training cohort and discrimination in the independent evaluation cohort were quantified using the ROC-AUC, with 95% confidence intervals (CIs) calculated using the DeLong method [45,46]. To account for class imbalance, precision-recall curves and the area under the precision-recall curve (PR-AUC) were additionally calculated for the overall training and evaluation cohorts [47]. The apparent training ROC-AUC was further corrected for optimism using the bootstrap procedure described above. As sensitivity analyses, ROC-AUCs with 95% CIs were also estimated within histologically restricted subsets of the training cohort, including simple carcinomas, grade 2–3 tumors, and grade 3 tumors. For adjusted analyses, we performed Firth’s penalized likelihood logistic regression because of the limited number of positive cases and the potential instability of estimates due to separation [48,49]. Odds ratios were obtained from the fitted regression coefficients, with 95% CIs based on profile penalized likelihood and *P* values derived from penalized likelihood-ratio tests. The RS was modeled as a continuous variable, and a series of models was fitted in which the association of RS with the outcome was adjusted for prespecified clinicopathologic covariates, individually or in combination. In the training cohort, the covariates included histologic grade, estrogen receptor (ER) status, neuter status, and age; in the evaluation cohort, they included histologic grade, clinical T stage, and age. Histologic grade was dichotomized as grade 3 versus grades 1–2 in the training cohort and grade 3 versus grade 2 in the evaluation cohort. Clinical T stage was modeled as a categorical variable. Analyses in the training cohort were restricted to cases with complete data for all covariates to allow comparison across model specifications. Adjusted odds ratios and 95% CIs for RS were reported across the different model specifications.

All analyses were performed in R version 4.4.3. Key R packages included GSVA (version 2.0.7), glmnet (version 4.1.10), logistf (version 1.26.1), pROC (version 1.19.0.1), and precrec (version 0.14.5).

## Results

### Cohort characteristics and outcome distributions

The training cohort included 113 dogs with CMC, of which 14 were classified as Spread-positive and 99 as Spread-negative (Table 1). All dogs were female. There was no evidence of a difference in age (*P* = 0.432) or neuter status (*P* = 0.497) between the two groups. In contrast, the distribution of histologic subtypes differed significantly between groups (*P* = 3.19 × 10^-9^). All Spread-positive tumors were simple carcinomas, comprising solid (78.6%), tubulopapillary (14.3%), and anaplastic (7.1%) subtypes. Spread-negative tumors were more heterogeneous, including tubulopapillary simple carcinoma (41.4%), complex carcinoma (36.4%), carcinoma in benign mixed tumor (14.1%), solid simple carcinoma (7.1%), and unannotated simple carcinoma (1.0%). The histologic grade also differed significantly according to lymphatic spread status (*P* = 1.59 × 10^-10^). All Spread-positive tumors were grade 2 or 3, including grade 3 in 92.9% and grade 2 in 7.1%, whereas Spread-negative tumors comprised 69.7% grade 1, 20.2% grade 2, and 10.1% grade 3 tumors. ER status also differed significantly between the groups (*P* = 1.87 × 10^-5^). Among tumors with available ER status, 57.1% of Spread-positive tumors were ER-positive and 42.9% were ER-negative, whereas 9.2% of Spread-negative tumors were ER-positive and 90.8% were ER-negative.

**Table 1.** Clinicopathologic characteristics of the 113 dogs in the training cohort stratified by lymphatic spread status.

| Variable | Spread<br>-negative (%) (N = 99) | Spread<br>-positive (%) (N = 14) | P value |
| --- | --- | --- | --- |
| **Age group (years)** |  |  | 0.432 |
| ≤ 12 | 48/94 (51.1) | 5/14 (35.7) |  |
| > 12 | 46/94 (48.9) | 9/14 (64.3) |  |
| Unknown | 5 | 0 |  |
| **Sex** |  |  | — |
| Female | 99/99 (100) | 14/14 (100) |  |
| **Neuter status** |  |  | 0.497 |
| Neutered | 35/96 (36.5) | 7/14 (50) |  |
| Intact | 61/96 (63.5) | 7/14 (50) |  |
| Unknown | 3 | 0 |  |
| **Histologic subtype** |  |  | 3.19e-09 |
| Simple carcinoma (Tubulopapillary) | 41/99 (41.4) | 2/14 (14.3) |  |
| Simple carcinoma (Solid) | 7/99 (7.1) | 11/14 (78.6) |  |
| Simple carcinoma (Anaplastic) | 0/99 (0) | 1/14 (7.1) |  |
| Simple carcinoma (Unannotated) | 1/99 (1) | 0/14 (0) |  |
| Carcinoma in benign mixed tumor | 14/99 (14.1) | 0/14 (0) |  |
| Complex carcinoma | 36/99 (36.4) | 0/14 (0) |  |
| **Histologic grade** |  |  | 1.59e-10 |
| 1 | 69/99 (69.7) | 0/14 (0) |  |
| 2 | 20/99 (20.2) | 1/14 (7.1) |  |
| 3 | 10/99 (10.1) | 13/14 (92.9) |  |
| **Estrogen receptor status** |  |  | 1.87e-05 |
| Negative | 89/98 (90.8) | 6/14 (42.9) |  |
| Positive | 9/98 (9.2) | 8/14 (57.1) |  |
| Unknown | 1 | 0 |  |

Values are presented as n/N (%) where N excludes missing or unknown data. Spread positivity was defined by the presence of tumor cells in peritumoral lymphatic vessels. *P* values were calculated using Pearson's chi-square test or Fisher's exact test, as appropriate (Fisher's exact test was used when expected cell counts were < 5). No *P* value was calculated for sex because there was no variation across groups.

The evaluation cohort included 27 dogs with CMC, comprising 13 LN-positive (48.1%) and 14 LN-negative (51.9%) cases (Table 2). All cases were classified as simple carcinomas and were ER-negative. Grade 3 tumors accounted for 53.8% and 64.3% of LN-positive and LN-negative cases, respectively, with the remaining cases classified as grade 2. Clinical T stage distributions varied between the groups. Among LN-positive tumors, 46.2% were T1 and 53.8% were T2, with no T3 tumors. For LN-negative tumors, 7.1% were T1, 64.3% were T2, and 28.6% were T3.

**Table 2.**
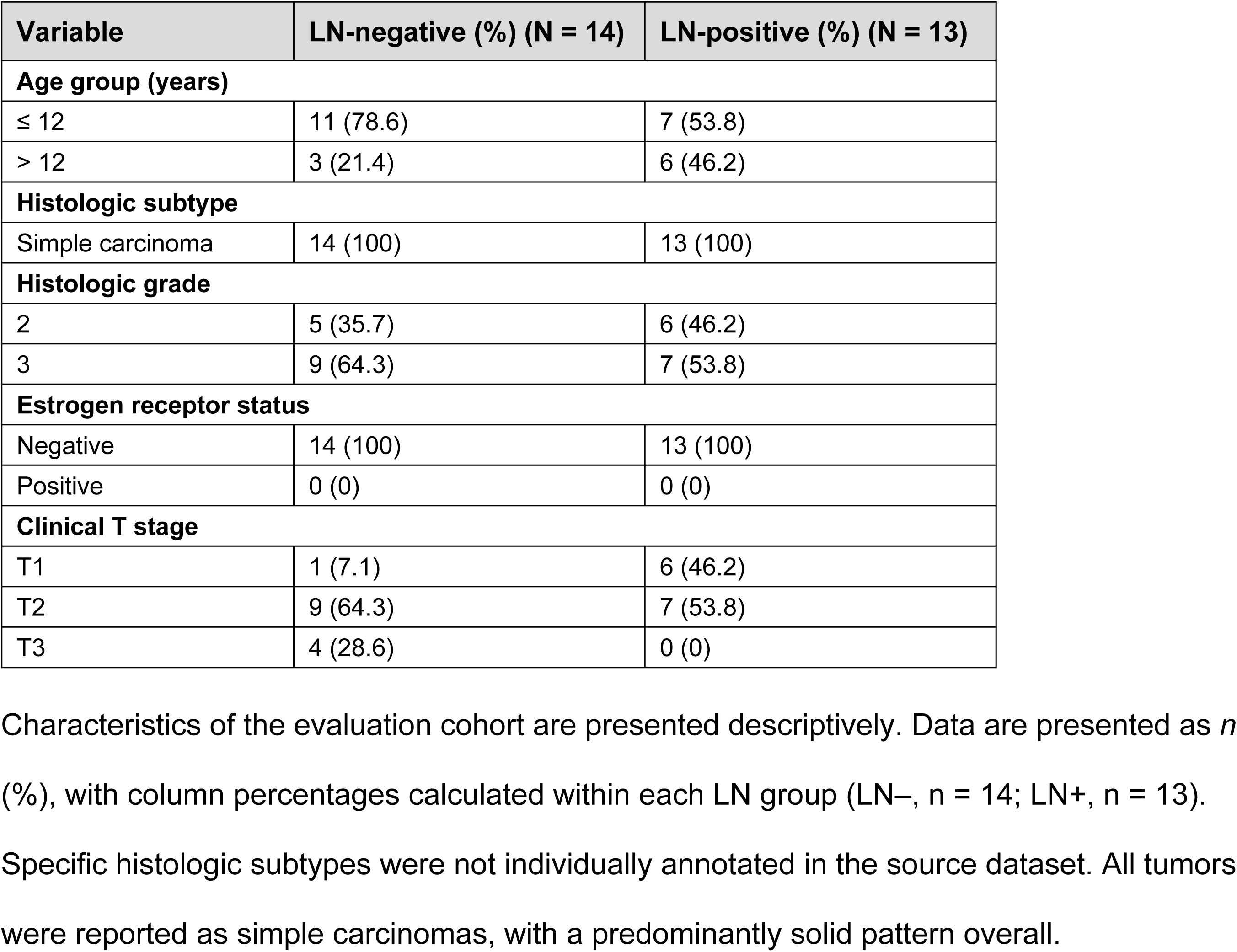
Clinicopathologic characteristics of the 27 dogs in the evaluation cohort stratified by LN metastasis status.

### Development of the lymphatic spread-associated transcriptomic risk score

Literature-guided curation yielded seven candidate gene sets for LASSO logistic regression, all of which showed significant univariable associations with lymphatic spread status in the training cohort (Table 3).

**Table 3.**
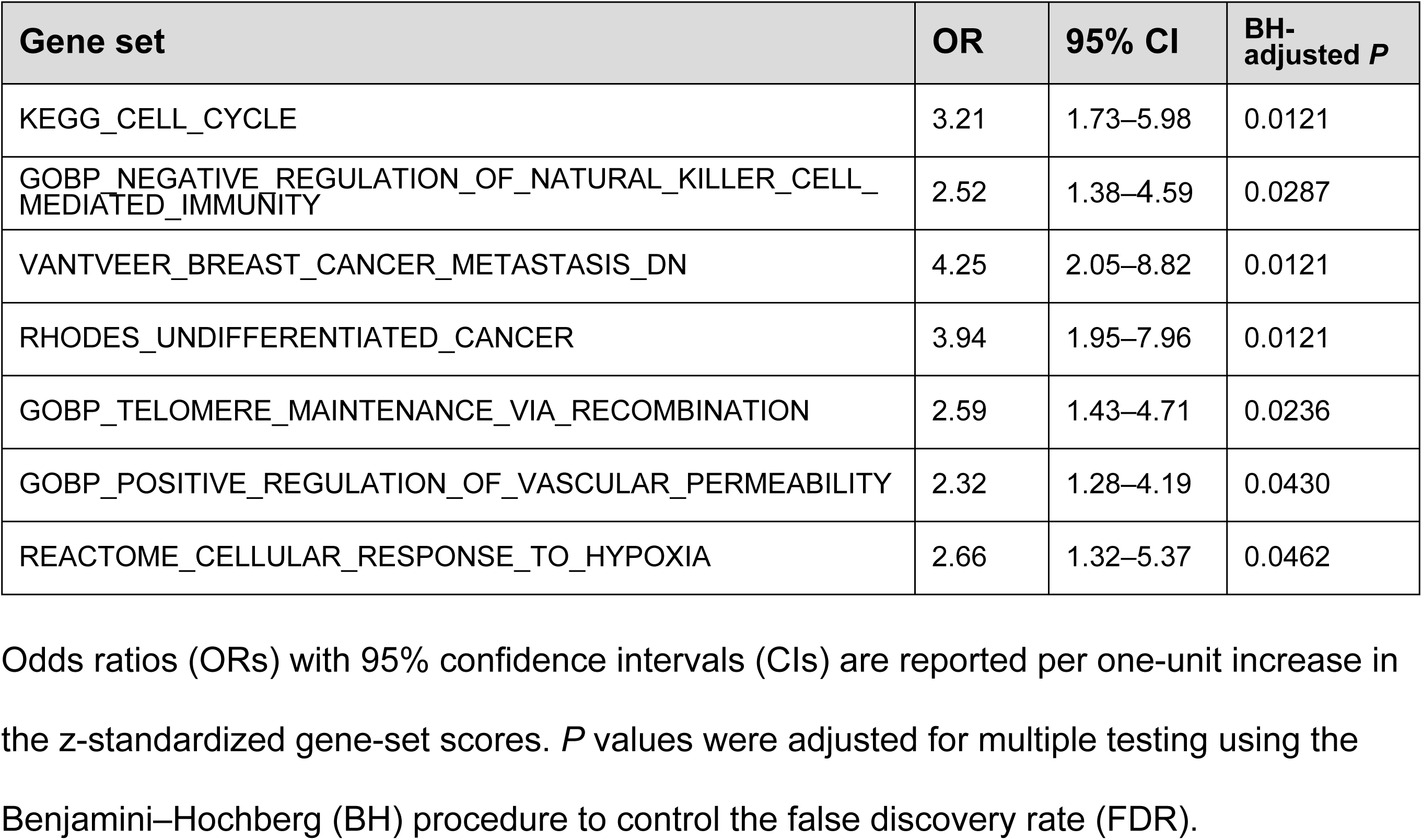
Univariable logistic regression results for the seven candidate gene sets selected for LASSO modeling.

LASSO logistic regression was performed using the seven candidate gene sets. The final model was fitted to the full training cohort using the median λ1SE obtained across 100 repetitions of five-fold cross-validation (S3 Fig). For visualization, the representative run shown in Fig 1 was selected as the repetition whose λ1SE was closest to the median λ1SE on the -log(λ) scale.

**Fig 1.**
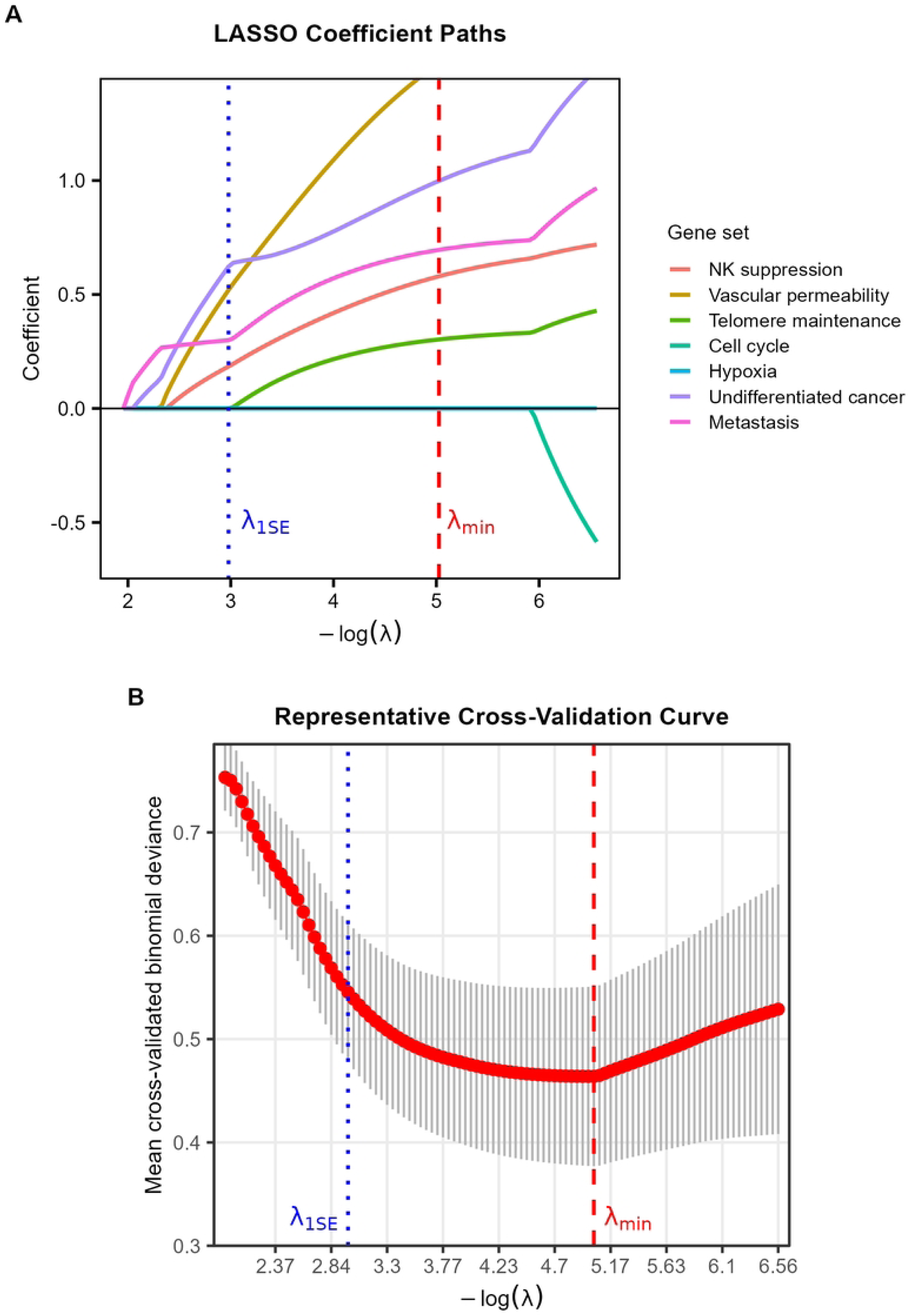
Repeated cross-validation and LASSO selection of gene-set features. **(A)** LASSO coefficient paths for the seven candidate gene sets as a function of - log(λ) in the representative cross-validation run. The blue dotted line indicates λ1SE, the largest λ with cross-validated binomial deviance within one standard error of the minimum, and the red dashed line indicates λmin, the λ yielding the minimum cross-validated deviance. **(B)** Five-fold cross-validation curve from the representative run, showing the mean cross-validated binomial deviance ±1 standard error. The blue dotted line and the red dashed line indicate λ1SE and λmin, respectively.

Four gene sets were retained in the final model: RHODES_UNDIFFERENTIATED_CANCER, GOBP_POSITIVE_REGULATION_OF_VASCULAR_PERMEABILITY, VANTVEER_BREAST_CANCER_METASTASIS_DN, and GOBP_NEGATIVE_REGULATION_OF_NATURAL_KILLER_CELL_MEDIATED_IM MUNITY. The four gene sets retained in the final model were also selected with relatively high frequencies across bootstrap resamples (Table 4). The composition of the four retained gene sets after common-gene restriction is provided in S4 Table.

**Table 4.**
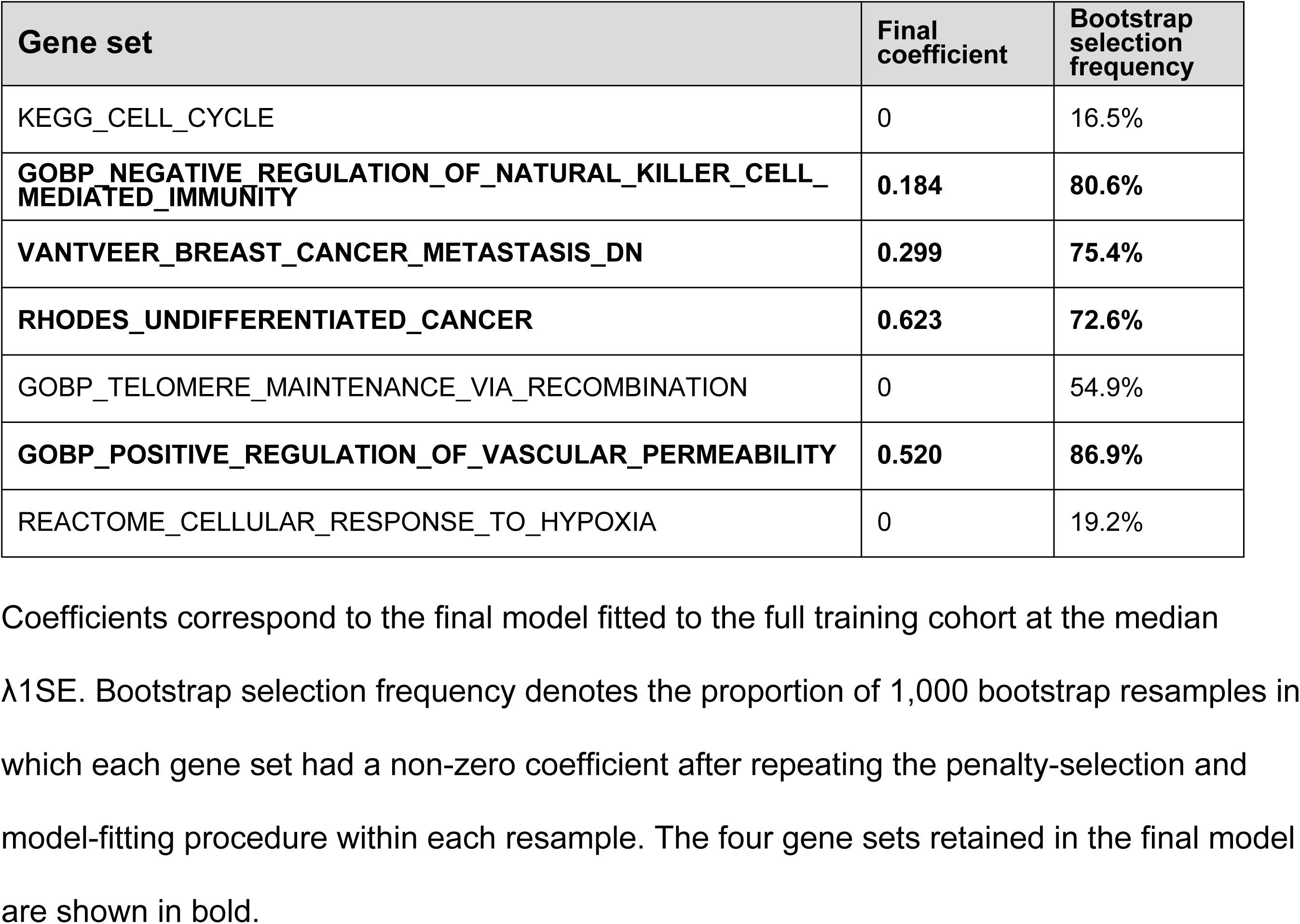
Final model coefficients and bootstrap selection frequencies of the seven candidate gene sets.

| Gene set | Final coefficient | Bootstrap selection frequency |
| --- | --- | --- |
| KEGG_CELL_CYCLE | 0 | 16.5% |
| **GOBP_NEGATIVE_REGULATION_OF_NATURAL_KILLER_CELL_MEDIATED_IMMUNITY** | **0.184** | **80.6%** |
| **VANTVEER_BREAST_CANCER_METASTASIS_DN** | **0.299** | **75.4%** |
| **RHODES_UNDIFFERENTIATED_CANCER** | **0.623** | **72.6%** |
| GOBP_TELOMERE_MAINTENANCE_VIA_RECOMBINATION | 0 | 54.9% |
| **GOBP_POSITIVE_REGULATION_OF_VASCULAR_PERMEABILITY** | **0.520** | **86.9%** |
| REACTOME_CELLULAR_RESPONSE_TO_HYPOXIA | 0 | 19.2% |

Coefficients correspond to the final model fitted to the full training cohort at the median λ1SE. Bootstrap selection frequency denotes the proportion of 1,000 bootstrap resamples in which each gene set had a non-zero coefficient after repeating the penalty-selection and model-fitting procedure within each resample. The four gene sets retained in the final model are shown in bold.

The final transcriptomic risk score was calculated as follows:

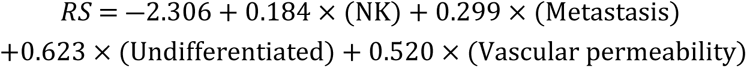

(where NK [natural killer], Metastasis, Undifferentiated, and Vascular permeability denote the standardized ssGSEA scores for GOBP_NEGATIVE_REGULATION_OF_NATURAL_KILLER_CELL_MEDIATED_IMMUNITY, VANTVEER_BREAST_CANCER_METASTASIS_DN, RHODES_UNDIFFERENTIATED_CANCER, and GOBP_POSITIVE_REGULATION_OF_VASCULAR_PERMEABILITY, respectively.)

The coefficients estimated in the training cohort were applied to the evaluation cohort without model refitting.

### Risk score distribution and sensitivity analyses

The RS was significantly higher in Spread-positive than in Spread-negative tumors in the overall training cohort (*P* = 5.80 × 10^-8^; Fig 2A). This difference remained significant in sensitivity analyses restricted to simple carcinomas (BH-adjusted *P* = 1.67 × 10^-7^), grade 2–3 tumors (BH-adjusted *P* = 1.10 × 10^-4^), and grade 3 tumors (BH-adjusted *P* = 0.030). Similarly, in the evaluation cohort, RS was significantly higher in LN-positive than in LN-negative tumors both overall (*P* = 2.59 × 10^-4^; Fig 2B) and among grade 3 tumors (*P* = 0.016).

**Fig 2.**
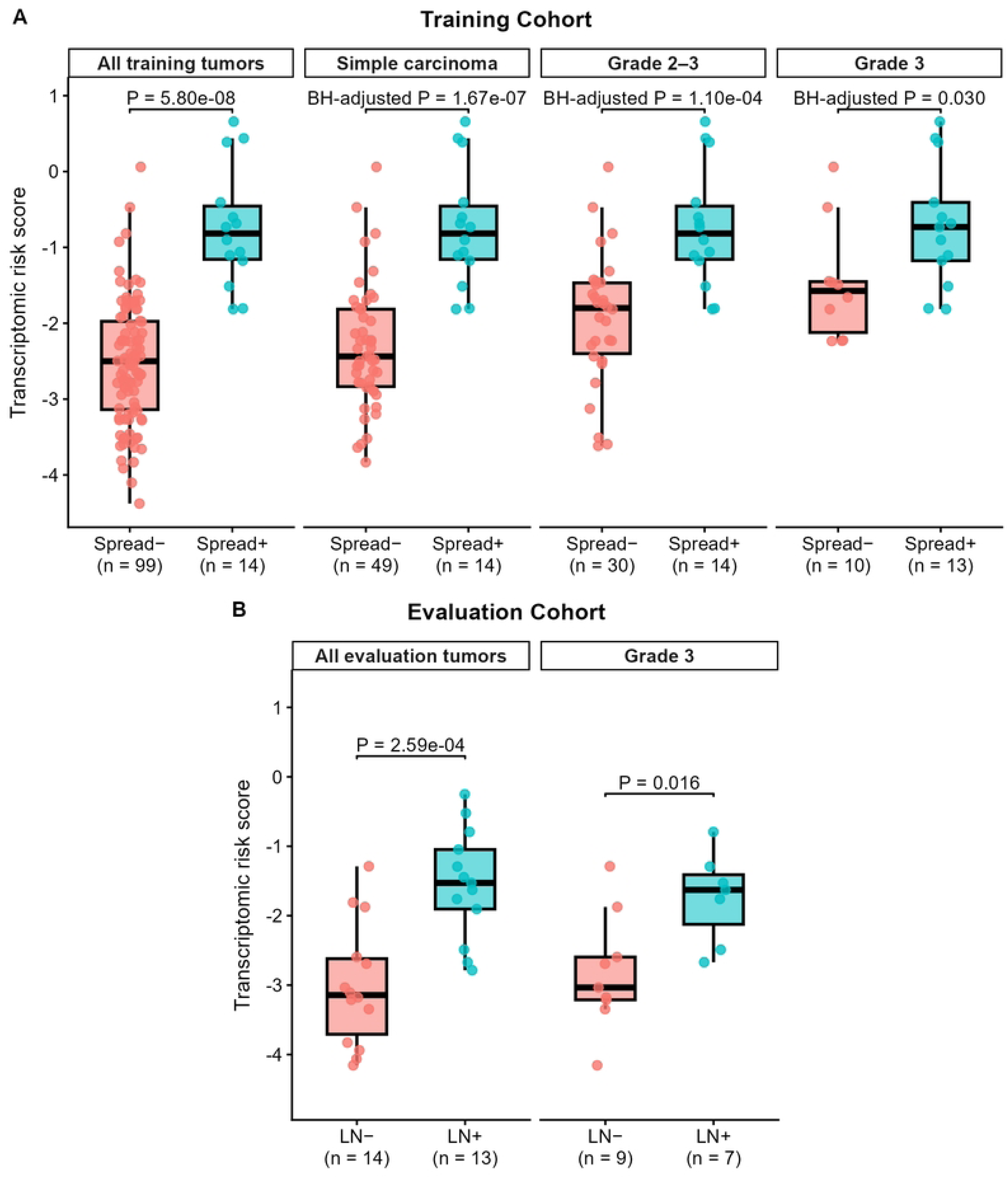
Distribution of transcriptomic risk scores according to locoregional dissemination status and sensitivity analyses in histologically restricted subsets. **(A)** Transcriptomic risk score according to lymphatic spread status in the training cohort, shown for all tumors and in sensitivity analyses restricted to simple carcinomas, grade 2–3 tumors, and grade 3 tumors. **(B)** RS according to LN metastasis status in the evaluation cohort, shown for all tumors and for the grade 3 subset. Individual points represent tumors, and boxplots show the median and interquartile range. Displayed *P* values are from two-sided Wilcoxon rank-sum tests; *P* values for the three training-cohort sensitivity analyses are adjusted using the Benjamini–Hochberg method.

### Discriminative performance of the risk score

In the training cohort, the RS showed strong apparent discrimination for lymphatic spread, with an ROC-AUC of 0.949 (95% CI, 0.906–0.993) and a PR-AUC of 0.725 (Fig 3A,B). Bootstrap internal validation estimated a mean optimism of 0.029, yielding an optimism-corrected ROC-AUC of 0.920. Discrimination was attenuated but remained above chance in histologically restricted sensitivity analyses, with ROC-AUCs of 0.926 (95% CI, 0.862–0.989) among simple carcinomas, 0.855 (95% CI, 0.739–0.971) among grade 2–3 tumors, and 0.769 (95% CI, 0.560–0.979) among grade 3 tumors (S2 Fig). In the independent evaluation cohort, the RS showed an ROC-AUC of 0.890 (95% CI, 0.766–1.000) and a PR-AUC of 0.869 (Fig 3C,D).

**Fig 3.**
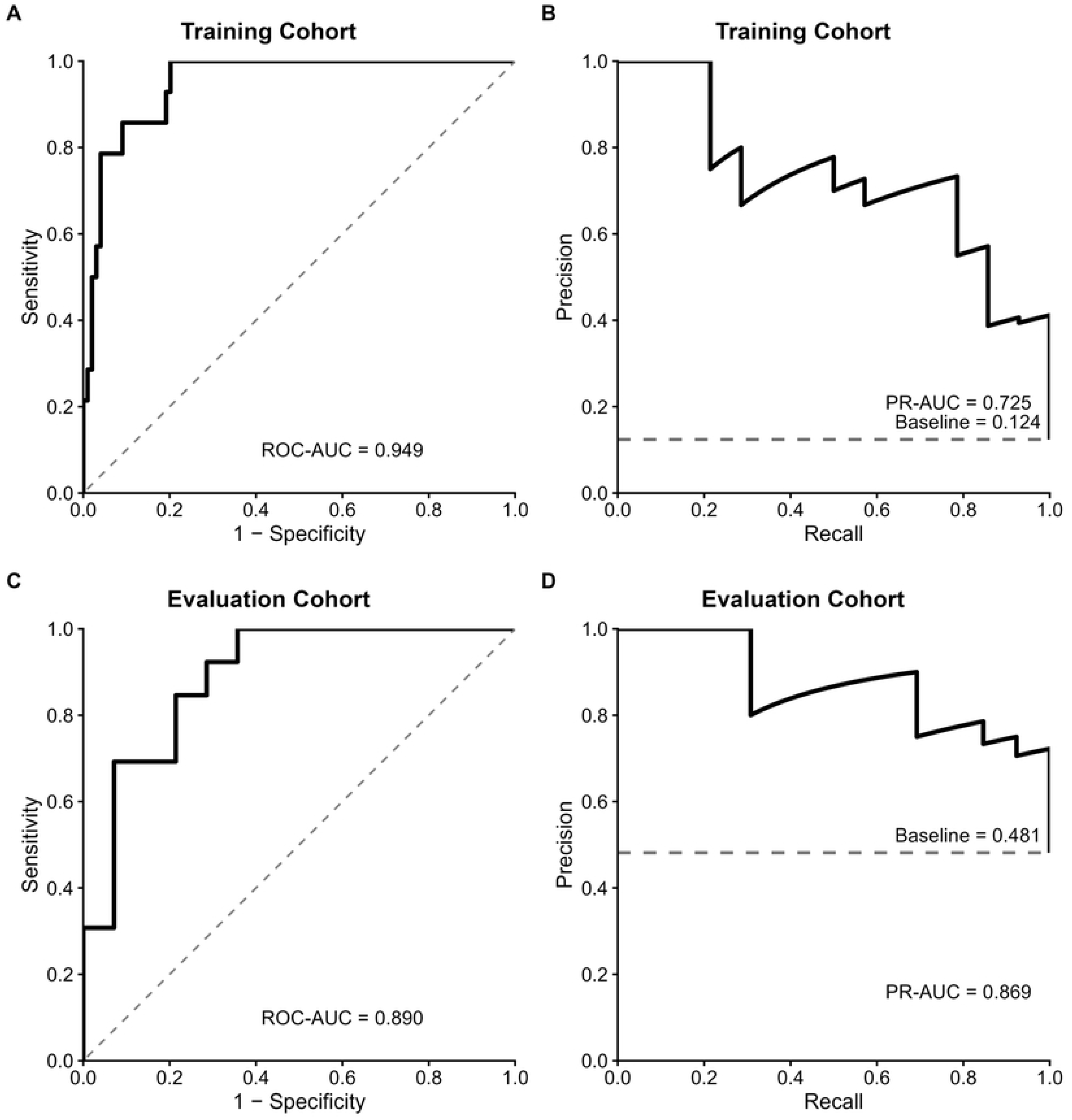
ROC and PR curves for the training cohort (lymphatic spread outcome) and evaluation cohort (nodal metastasis outcome). (A), **(C)** Receiver operating characteristic (ROC) curves and area under the curve (AUC) values in the training cohort and evaluation cohort. **(B), (D)** Precision-recall (PR) curves and PR-AUC values in the training cohort and evaluation cohort.

### Association of RS with locoregional dissemination across clinicopathologic adjustment models

Firth logistic regression analyses were used to assess the robustness of the association between continuous RS and locoregional dissemination across different clinicopathologic adjustment models (Table 5). Odds ratios for RS are reported per one-unit increase in the score. In the training cohort, among the 107 cases with complete covariate data, higher RS remained significantly associated with lymphatic spread across models adjusted for histologic grade, ER status, neuter status, and age, individually or in combination. Similarly, in the evaluation cohort, the association between higher RS and LN metastasis was retained across models incorporating histologic grade, clinical T stage, and age.

**Table 5.** Association of the transcriptomic risk score with locoregional dissemination across clinicopathologic adjustment models.

| Population and model | N (events) | OR (95% CI) | P value |
| --- | --- | --- | --- |
| Training cohort — Outcome: lymphatic spread |  |  |  |
| Unadjusted | 107 (14) | 11.11 (4.17–45.29) | 2.55e-10 |
| + Histologic grade (3 vs 1–2) |  | 4.29 (1.38–19.76) | 0.009 |
| + Estrogen receptor status |  | 9.78 (3.65–39.97) | 1.46e-08 |
| + Neuter status |  | 11.68 (4.18–50.57) | 3.80e-10 |
| + Age |  | 9.51 (3.67–37.30) | 3.56e-09 |
| All available covariates |  | 4.42 (1.34–22.91) | 0.011 |
| Evaluation cohort — Outcome: LN metastasis |  |  |  |
| Unadjusted | 27 (13) | 5.24 (1.92–23.13) | 2.98e-04 |
| + Histologic grade (3 vs 2) |  | 4.82 (1.83–19.81) | 4.12e-04 |
| + Clinical T stage |  | 3.96 (1.37–18.75) | 0.008 |
| + Age |  | 5.59 (1.87–29.79) | 6.70e-04 |
| All available covariates |  | 3.51 (1.21–17.32) | 0.019 |

Firth's penalized likelihood logistic regression was used to estimate the association between the continuous RS and lymphatic spread in the training cohort and LN metastasis in the evaluation cohort. Odds ratios (ORs) and 95% confidence intervals (CIs) are reported per one-unit increase in RS. Each adjusted model included RS and the indicated clinicopathologic covariate; the "All available covariates" model included all listed covariates simultaneously. N (events) indicates the number of tumors included in the analysis and the number with the corresponding positive outcome, respectively.

## Discussion

### Transcriptomic features associated with locoregional dissemination

In this study, we developed a primary-tumor transcriptome-derived risk score associated with locoregional dissemination in CMC and evaluated its association with histopathologically defined nodal metastasis in an independent cohort. Previous molecular studies have demonstrated that metastatic behavior in CMC is accompanied by changes in primary-tumor gene expression, miRNA profiles, and individual candidate transcripts [23–25]. The present findings extend this body of work by integrating dissemination-associated transcriptional features at the gene-set level into a sparse RS and evaluating the resulting signal across independent cohorts generated using different transcriptomic platforms.

In the training cohort, histologic grade and subtype differed markedly according to lymphatic spread status, and the lower ROC-AUC estimates observed after histologic restriction were consistent with partial overlap between the transcriptomic signal captured by the RS and conventional histopathologic features of tumor aggressiveness, although the small subgroup sizes limit this interpretation.

ER status also differed markedly according to spread status, with ER positivity being more frequent among Spread-positive tumors. This pattern contrasts with the generally reported association between reduced ER expression and more aggressive canine mammary tumor phenotypes [50,51]. Given the small number of Spread-positive tumors and the marked clinicopathologic imbalance between groups, this unexpected association should be interpreted cautiously.

Nevertheless, the association between RS and lymphatic spread remained significant after multivariable adjustment including histologic grade and ER status, and the RS remained discriminatory in the independent evaluation cohort, in which all tumors were ER-negative. Thus, although the RS appears to capture transcriptional features that overlap with established indicators of tumor aggressiveness, its association with lymphatic spread was not solely attributable to these clinicopathologic characteristics.

### Biological interpretation of the risk score

The four gene-set-level features retained in the final RS provide biological context for the primary-tumor transcriptomic state associated with locoregional dissemination. Because the RS was derived from bulk transcriptomic data, these features should be interpreted as coordinated gene-set-level summaries rather than direct evidence of specific causal mechanisms, cellular abundance, or functional immune activity. Two retained features were empirical cancer-related expression signatures. RHODES_UNDIFFERENTIATED_CANCER represents genes enriched in undifferentiated cancers [52], suggesting that locoregional dissemination may be associated with a less differentiated primary-tumor state. The retention of VANTVEER_BREAST_CANCER_METASTASIS_DN, derived from the original breast cancer outcome signature [20], is also consistent with the concept that clinically relevant dissemination-related information can be reflected in primary-tumor expression profiles.

The remaining features were curated biological-process gene sets related to vascular permeability and natural killer (NK) cell-mediated immunity. The vascular permeability-related feature may reflect vascular and stromal remodeling associated with aggressive tumor behavior. In canine mammary tumors, higher vascular endothelial growth factor (VEGF) protein immunoreactivity was associated with malignancy, poorer differentiation, and greater microvessel density [53]. In human breast cancer, *VEGFA* mRNA was highly expressed in ductal carcinoma cells, with receptor transcripts localized to adjacent endothelial cells [54]. Together with the established role of VEGF-A in vascular hyperpermeability, angiogenesis, and stromal formation [55], these findings support the biological relevance of this feature to a dissemination-associated tumor microenvironment.

The NK cell-related feature may reflect attenuation of innate immune surveillance. Impaired NK-cell phenotype and function have been reported in invasive human breast cancer [56], and NK cells are recognized contributors to metastatic immunosurveillance [57,58]. Thus, the enrichment of negative regulation of NK cell-mediated immunity may represent an immune-evasive component of the dissemination-associated primary-tumor state.

Taken together, the RS should be viewed as a composite index integrating reduced differentiation, metastasis-related expression programs, vascular or stromal remodeling, and altered innate immune surveillance, rather than as a direct measure of a single metastatic pathway.

### Limitations

This study has several limitations. First, the number of positive cases was limited in both cohorts, reducing the precision of effect estimates and constraining the complexity of clinicopathologic adjustment. Firth’s penalized likelihood logistic regression was therefore used to mitigate instability related to sparse data and separation. The small sample sizes of the histologically restricted subgroups and their wide confidence intervals limit the precision of the subgroup discrimination estimates.

Second, the training and evaluation cohorts were generated using different transcriptomic platforms. Although gene-set membership was restricted to genes represented in both datasets and gene-set-level scores were used to reduce platform-specific effects, residual cross-platform variation cannot be excluded.

Moreover, ssGSEA scores were standardized within each cohort, facilitating relative discrimination within cohorts but limiting direct comparison of absolute RS values and straightforward transfer of a fixed threshold across platforms.

Third, bootstrap internal validation repeated penalty selection and model fitting conditional on the seven curated candidate gene sets and therefore did not account for uncertainty introduced by the preceding gene-set screening and literature-guided curation; residual optimism from these upstream steps cannot be excluded.

Fourth, the outcome definitions differed between cohorts. The training endpoint reflected peritumoral lymphatic invasion, whereas the evaluation endpoint was histopathologically confirmed lymph node metastasis. Although both endpoints represent locoregional dissemination, they may reflect different stages and manifestations of metastatic progression.

Finally, both endpoints represented locoregional dissemination status at the time of surgical resection rather than prospectively observed metastatic progression.

Accordingly, the RS should be interpreted as a marker of a dissemination-associated primary-tumor state rather than as a predictor of future metastasis.

## Conclusions

In conclusion, this study developed a transcriptomic RS associated with locoregional dissemination in CMC and evaluated its performance in an independent cohort with histopathologically confirmed LN metastasis. The RS may represent a complementary molecular layer for identifying dissemination-associated primary-tumor biology, but should not yet be considered a clinically validated prognostic or nodal-staging tool. Prospective evaluation of a prespecified score in independent cohorts with standardized pathological and longitudinal outcome assessment is required to establish its clinical utility and prognostic significance.

## Data availability statement

The transcriptomic datasets analyzed in this study are publicly available in the GEO under accession numbers GSE119810 and GSE20718. The processed data and supplementary results generated during the present analyses are included in this article and its Supplementary Information. The principal analysis scripts used in this study are publicly available at https://github.com/brianjingo420-max/riskscoreGeneration.

## Supporting information

**S1 Fig. Pairwise Spearman correlation matrix of the seven candidate gene-set ssGSEA scores in the training cohort.** Values in each cell represent Spearman’s ρ. The lower triangle is shown for clarity, with red indicating positive and blue indicating negative correlations. NK, natural killer; Vasc perm, vascular permeability; Metastasis (vV), van’t Veer breast cancer metastasis signature.

**S2 Fig. Discrimination of the transcriptomic risk score in histologically restricted subsets of the training cohort.** ROC-AUC estimates and 95% confidence intervals are shown for the overall training cohort (n = 113; events = 14), simple carcinoma (n = 63; events = 14), grade 2–3 tumors (n = 44; events = 14), and grade 3 tumors (n = 23; events = 13). Points indicate ROC-AUC estimates and horizontal bars indicate 95% confidence intervals calculated using the DeLong method.

**S3 Fig. Distribution of -log(λ1SE) values obtained across 100 repeated five-fold cross-validation runs.** The dashed vertical line indicates the -log of the median λ1SE across repetitions, which was used to define the regularization level for the final model.

**S1 Table. Full results of univariable gene-set screening associated with lymphatic spread in the training cohort.** Univariable logistic regression was performed for each gene set using standardized single-sample gene set enrichment analysis scores in the training cohort. *P* values were adjusted for multiple testing using the Benjamini–Hochberg method. Gene sets with false discovery rate < 0.05 are shown. OR, odds ratio; CI, confidence interval; SE, standard error; FDR, false discovery rate; BH, Benjamini–Hochberg.

**S2 Table. Literature-guided selection of representative gene sets for candidate biological domains.** Seven biological domains were defined through literature-guided curation of gene sets meeting the prespecified univariable screening criterion. One representative gene set was selected from each domain for inclusion in least absolute shrinkage and selection operator logistic regression.

**S3 Table. Representative gene sets considered during literature-guided curation within each biological domain.** Gene sets considered during literature-guided curation are shown by biological domain. “Selected” denotes the representative gene set retained for subsequent least absolute shrinkage and selection operator modeling, whereas “Alternative” denotes other biologically related gene sets considered within the same domain.

**S4 Table. Characteristics and matched gene membership of the four gene sets retained in the final risk score.** Gene-set descriptions and original membership were obtained from the Molecular Signatures Database. Genes represented in both the training and evaluation datasets after ortholog mapping and preprocessing were used for single-sample gene set enrichment analysis.

**S5 Table. λ1SE values obtained across 100 repeated five-fold cross-validation runs.** The one-standard-error penalty parameter (λ1SE) obtained from each repetition is listed in ascending order. The median λ1SE across 100 repetitions was used to fit the final least absolute shrinkage and selection operator logistic regression model in the full training cohort. The corresponding distribution of λ1SE values is shown in S3 Fig.

**S6 Table. Detailed results of the Firth penalized logistic regression models summarized in Table 5**. All fitted model terms except intercepts are shown. CI, confidence interval; ER, estrogen receptor; OR, odds ratio; RS, risk score; SE, standard error.

**S7 Table. Individual-level data underlying the risk-score distributions in Fig 2 and the ROC and precision-recall curve analyses in Fig 3**. Outcome was coded as 1 for the presence and 0 for the absence of lymphatic spread in the training cohort or LN metastasis in the evaluation cohort. The simple-carcinoma and histologic-grade variables define the subgroup analyses presented in Fig 2.

**S8 Table. Individual-level standardized scores for the seven candidate gene sets used in the correlation analysis presented in S1 Fig.** Each row represents an individual tumor from the training cohort. Gene-set scores were calculated using single-sample gene set enrichment analysis (ssGSEA) and z-standardized within the training cohort. These values were used to calculate the pairwise Spearman correlation coefficients shown in S1 Fig.

